# Detection of A-to-I microRNA editing with miRmedon reveals widespread co-editing of mature microRNA in the human brain

**DOI:** 10.1101/2022.12.23.521716

**Authors:** Amitai Mordechai, Alal Eran

## Abstract

microRNA (miRNA), key regulators of gene expression, are prime targets for adenosine deaminase acting on RNA (ADAR) enzymes. Although ADAR-mediated adenosine-to-inosine (A-to-I) miRNA editing has been shown to be essential for orchestrating complex processes, including neurodevelopment and cancer progression, only few human miRNA editing sites have been reported. Several computational approaches have been developed for the detection of miRNA editing in small RNAseq data, all based on the identification of systematic mismatches of ‘G’ at primary adenosine sites in known miRNA sequences. However, these methods have several limitations, including their ability to detect only one editing site per sequence (although editing of multiple sites per miRNA has been reproducibly validated), their focus on uniquely mapping reads (even though 20% of human miRNA are transcribed from multiple loci), and their inability to detect editing in miRNA harboring genomic variants (though 73% of human miRNA loci include a reported SNP or indel). To overcome these limitations, we developed miRmedon, that leverages large-scale human variation data, a combination of local and global alignments, and a comparison of the inferred editing and error distributions, for confident detection of miRNA editing in small RNAseq data. We demonstrate the improved performance of miRmedon as compared to currently available methods and describe its advantages. We further use miRmedon to discover editing haplotypes of mature human brain miRNA for the first time. We find that doubly edited mature miRNA are common in the adult human prefrontal cortex, most include a frequently edited site within the miRNA seed region, and are predicted to maintain a stable pre-miRNA structure. These results suggest that co-editing of mature miRNA could enable efficient shifting of gene expression programs.

## INTRODUCTION

Owing to their double-stranded structure, miRNA are considered prime targets for A-to-I RNA editing, a site-specific deamination of adenosine to inosine carried out by ADAR enzymes. By efficiently altering miRNA biogenesis and/or targeting, A-to-I editing can induce global gene expression shifts (1). Accordingly, A-to-I miRNA editing has been shown to be essential for orchestrating neuronal development and cancer progression (2,3). However, despite their emerging role in human health and disease, very few human miRNA editing sites have been reported (4).

Several computational approaches have been developed for the detection of miRNA editing in small RNAseq data, all based on the identification of systematic mismatches of ‘G’ at primary adenosine sites in known miRNA sequences (5-11) (**Supplementary Table S1**). A major barrier in confidently detecting editing events in small RNAseq is cross-mapping, namely the likely mapping of a putatively-edited miRNA sequence to multiple loci, including those from which it did not originate (6). For this reason, the dominating miRNA editing detection approach restricts its analysis to uniquely mapping small RNA reads (11). However, this requirement neglects potential editing in 20.3% of all mature human miRNA that could have originated from several loci (12). A more tolerant approach was taken by Zheng et al. (10), using a cross-mapping correction method (6). Still, this approach cannot detect multiple editing sites per miRNA, since considering all possible editing events in mature miRNA aggravates the cross-mapping issue. Moreover, since all existing methods rely on alignment to the genome or miRBase with one mismatch, they are unable to detect editing in miRNA harboring genomic variants.

To overcome these limitations, we developed miRmedon, (<u>miR</u>NA <u>m</u>ultiple <u>ed</u>iting detecti<u>on</u>), a novel framework for the confident detection of multiple editing sites in a single read. miRmedon is based on a combination of local and global alignments to the genome and transcriptome, as well as consideration of large-scale population variation data, and a comparison of the inferred editing distribution to the sequencing error distribution. We demonstrate the improved performance of miRmedon for *de novo* detection of A-to-I miRNA editing, as compared to currently available methods, and use it to explore co-editing of mature miRNA in the adult human brain.

## MATERIALS AND METHODS

### Estimation of miRNA sequence complexity

Sequence complexity was estimated using Shannon’s entropy, as described in Equation 1.

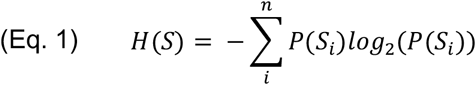

where *P(S*_*i*_) is the relative abundance of each nucleotide *i ∈* {*A, T, C, G*} in string *S*. Mature miRNA sequences with *H(S) < 1*.*4* were considered low complexity sequences. This threshold was determined based on the miRBase 21 H(S) distribution, following outlier removal.

### e-miRBase generation

In constructing an e-miRBase based on the 2,656 mature miRNA sequences of miRBase 21, we first discarded 59 low complexity sequences. Then, 11,421 putative editing sites were considered in all primary adenosine sites of the remaining 2,597 mature miRNAs, with the exception of those in two most 3’ positions, shown to be highly modified and a potential source of false positives (11). Of these, 2,060 potential sites were found to overlap with a gnomAD SNP or be located within 5bp of a gnomAD indel, and were therefore ignored to improve specificity. We note that future versions of miRmedon will enable tuning this filtration step via a user-determined minor allele frequency cutoff. Next, all possible combinations of sequences containing a guanosine in the remaining 9,361 primary adenosine sites were generated, and those resulting in low complexity sequences were removed, yielding 131,880 sequences that were globally aligned to the genome and transcriptome using SeqMap with no mismatches. Of these, 839 were fully aligned to the genome and/or transcriptome, and therefore excluded from the final e-miRBase index, which included 133,638 edited and unedited e-miRNAs, encompassing 9,209 possible editing sites.

While currently available miRNA editing detection approaches use dbSNP to identify genomic variants that could potentially be mistaken for editing events, miRmedon leverages gnomAD for this purpose. In comparison to dbSNP, whose credibility has been criticized by many (e.g., (13)), gnomAD provides a high confidence set of rigorously called variants from diverse human populations. Moreover, miRmedon considers genomic variants of all allele frequencies low or high, following the approach taken by all currently available miRNA editing detection pipelines (**Supplementary Table S1**). These and nearly all general RNA editing detection methods, including the popular REDItools (14) and RNAEditor (15), prefer high specificity and consider all genomic variants independent of their allele frequency, at the cost of potentially increasing false negatives.

### Alignment

miRmedon uses a combination of local and global alignments. First, to align the e-miRBase collection to the human genome and transcriptome, a global aligner such as Bowtie (16) or SeqMap (17), is used. Bowtie is used with -a --best, -S and -v options, while SeqMap is used with the /output_all_matches option. Notably, Bowtie and SeqMap with the above-mentioned parameters yield the exact same results.

RNAseq reads are locally aligned to e-miRBase using STAR (18), with the following parameters:

--alignIntronMin 1
--outFilterMultimapNmax 200
--outFilterMatchNmin 12
--outFilterMismatchNoverLmax 0.08
--seedSearchStartLmax 6
--winAnchorMultimapNmax 2000
--outFilterMultimapScoreRange 0
--outSAMtype BAM Unsorted
--outFilterMismatchNmax 1
--outSAMprimaryFlag AllBestScore
--outSAMattributes NH AS NM MD
--outWigType None

Of note, STAR was selected over other commonly used local aligners, such as Bowtie2, Kallisto, and Salmon, owing to its demonstrated superior precision and recall for aligning short unspliced reads to short sequences (19).

### Reference-free quality-dependent consensus sequence inference

Following local alignment of reads to the e-miRBase collection and subsequent filtration, multiple sequence alignment (MSA) is applied to each set of reads *R*_*i*_ aligned to *e-miRNA*_*i*_ using Mafft (20,21), while utilizing the full length reads and the --op 10 option. For each aligned read *r*_*ij*_*∈ R*_*i*_, in each position *k*, a discrete probability function representing the probability of observing each of the calls in the set *C =* {*A, T, C, G, N, -*} is formed (with ‘-’ representing a gap as determined by the MSA). Provided that *r*_*i,j,k*_ is an unambiguous nucleotide (A, T, C or G), the probability *p* of the observed base is calculated from the Phred score 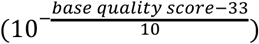 while the probability of observing each of the alternative nucleotides is (1-*p)/3*. The probability of observing an ambiguous character (N/-) is set to *ε*. On the other hand, if *r*_*i,j,k*_ = ‘N’ or ‘-’, the probability for an ambiguous character is set to *1 – ε*, and the probability to observe each of the legitimate nucleotides is *ε/4*. Then, the probability to observe the character *C*_*i,k*_ *∈ C*, which is used to calculate the merged probability function of observing *C*_*i,k*_ in position *k* based in all reads of cluster *Ri*, is shown in Equation 2 (Genest and Zidek, 1986). Finally, the character in each position *k* of the consensus sequence is determined to be *C*_*i,k*_ of the highest probability, as calculated by Equation 2. Positions with multiple equally likely characters are inferred as ‘N’.

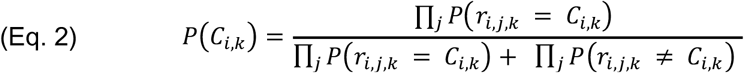

The inferred consensus sequences are then globally aligned to the reference human genome and transcriptome using SeqMap or Bowtie, as detailed above.

### Editing site inference

miRmedon uses Monte Carlo sampling to test whether an observed mismatch could be attributed to sequencing error. Specifically, for each putative editing site, two distributions are repeatedly sampled and compared: the posterior editing distribution and the sequencing error distribution. The first is a beta distribution with parameters α = 1 + the number of reads with G mapped to the site, and β = 1 + the number of reads with A mapped to the site; the second distribution is that formed by all base quality (Q) scores of the alternate base being examined. It is the distribution of 10^−Q/10^ scores of calls supporting the alternate base in the examined position, essentially representing the distribution of error probabilities of the observed alternate base among all its supporting reads.

Using repeated resampling (default 10,000 resamples), a random draw from the editing distribution is compared to a random draw from the sequencing error distribution of the alternate base. With the results of all resamples at hand, a P value is calculated as the fraction of resamples in which the sampled probability of sequencing error was greater than or equal to the sampled editing level. Thus, this P value represents the chance that the observed mismatch could be attributed to sequencing error. All calculated P values are corrected for multiple testing using the Benjamini-Hochberg method (22).

### Small RNAseq data

Publicly available small RNAseq data was downloaded from the NCBI Sequence Read Archive (SRA), using the accession numbers detailed in **Supplementary Table S2**.

### RNAseq quality control and pre-processing

Small RNA reads were first subject to quality control using FastQC (23). Unless already trimmed, adapters were removed using Cutadapt (24), followed by quality-based filtering with Trimmomatic (25), using parameters SLIDINGWINDOW:4:20 and MINLEN:15.

### miRmedon performance evaluation

miRmedon was compared to seven leading approaches for the detection of miRNA editing (**Supplementary Table S1**), of which the dominant one, namely that of Alon et al. (11), was used for benchmarking. Toward that goal, small RNAseq from pooled human brain, generated by Alon et al. (11), was used as input for miRmedon, using the default expression threshold of five reads and 100,000 resamples for P value estimation. In all, 6,047,953 reads that passed quality control were analysed by miRmedon. The detection rate was calculated as the percent of sites detected by miRmedon and/or the Alon et al. pipeline out of all sites that could possibly be detected based on miRNA expression in the examined sample. Sites that overlap with gnomAD variants (26) were not included in this calculation, thereby ignoring 12 previously reported sites, of which five were considered high-confidence sites by Pinto et al. (**Supplementary Tables S3** and **S4**).

### Classification of miRmedon-detected mismatches based on their similarity to high confidence A-to-I miRNA editing sites

A regularized logistic regression model was trained to classify miRmedon-detected mismatches based on their similarity to high-confidence A-to-I miRNA editing sites. The model’s features were the number of edited and unedited reads, the point estimate for the editing level based on a posterior beta distribution, its lower and upper 95% confidence interval bounds, the adjusted P value, the nucleotide immediately upstream to the site (represented by one hot encoding), the nucleotide immediately downstream to the site (similarly represented by one hot encoding), and a final binary feature indicating whether the site resides in the miRNA seed region. The first six features were scaled to Z scores and the model was trained on 15 high confidence A-to-I miRNA editing sites as true positives and 23 non A>G sites as true negatives, all detected in the pooled human brain sample when miRmedon was applied to detect all 12 possible substitutions. The model is detailed in **Supplementary Table S5** and its training set is detailed in **Supplementary Table S6**.

The model’s performance was evaluated using leave-one-out cross-validation (LOOCV). To assess miRmedon’s FDR, the model was then applied to 58 mismatches detected by applying miRmedon to each substitution separately. Specifically, since miRmedon directly detects A>G substitutions and performs multiple testing correction only on putative A-to-I editing sites, we replicated this process for each substitution. Accordingly, for each substitution, all sites with an adjusted P value < 0.05 for that substitution were included in the FDR assessment. We excluded from this analysis T>G mismatches at positions 12 and 15 of hsa-miR-92a-2-5p, which were detected only in doubly-edited reads with 100% editing, with no other form of this miRNA. Since it is highly implausible to have a miRNA edited in two adjacent sites with no evidence for transcripts edited in one or none of these sites, this is evidently a false positive. Notably, such anomaly was not detected elsewhere. The 58 sites included in the FDR assessment are detailed in **Supplementary Table S7**.

### Secondary structure analysis and visualizations

To examine the stability of co-edited miRNA, we used RNAfold of the ViennaRNA package version 2.4.14 (27) to predict the secondary structure and minimum free energy of the pre-miRNA containing inosine at both editing sites. Visualizations were made using RiboSketch (28).

### Detection of mature miRNA editing in the adult human brain

Small RNAseq data from 64 postmortem prefrontal cortex samples detailed in **Supplementary Table S2** were downloaded from the NCBI SRA, via accession GSE64977. Quality control and adapter trimming were performed as described above. miRmedon was used to detect miRNA editing sites using its default parameters detailed above, with the additional criterion of detection in at least three samples. For validation purposes, the expected number of edited reads was calculated using the raw read count of the non-edited miRNA and the editing level detected in the pooled human brain sample. Sites with at least five expected edited reads in at least three samples who lacked sufficient support for editing were deemed not validated. Otherwise, sites with insufficient coverage, as compared to the expected editing-supporting read count, were considered below the detection limit.

## RESULTS

### Description of miRmedon

#### e-miRBase generation

The first step of miRmedon includes the *in-silico* generation of an extended, human variation-aware, edited human miRNA collection, which we term e-miRBase (**Figure 1A**). For that purpose, miRBase (12) is used as a template for simulating all possible editing combinations, i.e. guanosines at primary adenosine sites, with the exception of the two most 3’ bases, which were shown to be highly modified and a potential source of false positives (11). Simulated editing sites mapped to a gnomAD SNP (26) or those within a user-defined distance from a gnomAD indel (default 5bp) are excluded. Additionally, low complexity sequences, including non-edited miRNAs, are discarded because of the high risk for spurious mappings of these sequences (**Materials and Methods**). The resulting e-miRNA sequences are then globally aligned to the full reference human genome (which includes not only chromosomes, but also scaffolds, patches, and haplotypes) and transcriptome, using SeqMap or Bowtie with no mismatches (16,17). e-miRNA sequences that perfectly match non-miRNA loci or transcripts are discarded.

**Figure 1.**
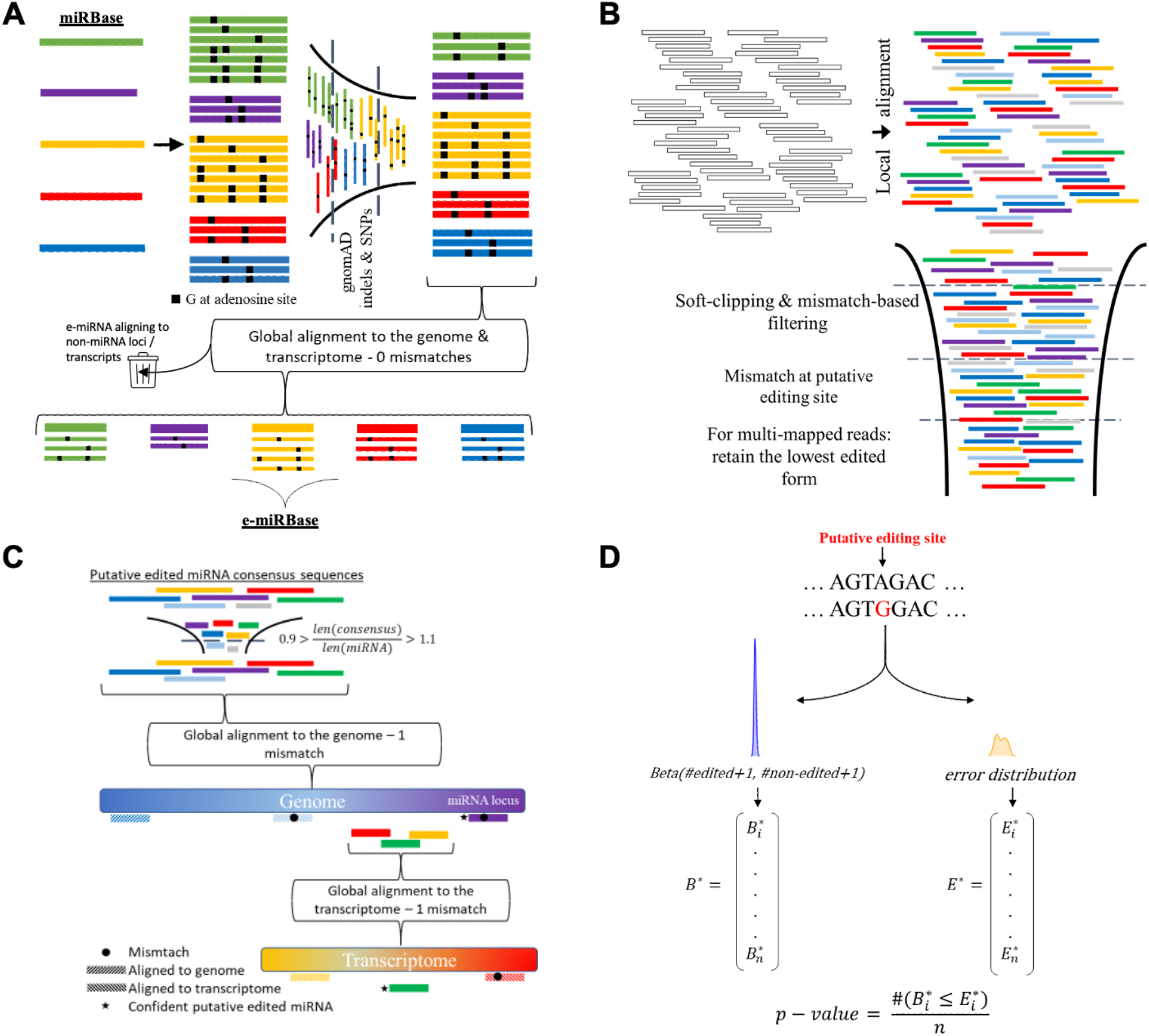
miRmedon framework. (**A**) e-miRBase generation. An extended version of miRBase is generated, in which each mature miRNA is represented by all possible combinations of putative editing sites, excluding those overlapping gnomAD variants or those at the 3’ ends. Each so-called e-miRNA sequence is then globally aligned to the full reference human genome and transcriptome, and e-miRNA aligning to non-miRNA loci or transcripts are discarded; (**B**) Read clustering. Small RNAseq reads are locally aligned to e-miRBase, and a series of alignment-based filters is employed; (**C**) Read classification. For each e-miRNA, multiple sequence alignment is performed followed by quality-dependent consensus sequence inference. After discarding consensus sequences of inappropriate length, the remaining consensus sequences are globally aligned to the reference human genome and transcriptome with 1 mismatch, and those mapping to non-miRNA loci are removed; (**D**) Editing site inference and quantification. For each putative editing site, editing levels are modeled as a beta distribution (B) based on the number and identity of reads mapped to the site. The error distribution (E) of a site is based on its base quality scores. A bootstrapped P value is calculated based on repeated resampling and comparison of the resulting distributions *B*^*\**^ and E^*\**^, as the number of times *B*^*\**^_*i*_ is less than or equal to *E*^*\**^_*i*_, divided by the number of bootstrap samples *n*.

#### Read clustering

The next miRmedon step revolves around local alignment of small RNAseq reads to the e-miRBase collection generated in the previous step (**Figure 1B**). Specifically, trimmed and quality inspected small RNAseq reads are aligned to e-miRBase using STAR (18). Several post-alignment filters and modifications are then employed. First, reverse complement alignments are discarded. Then, reads that were soft-clipped more than a user-defined threshold are removed (default 2). Similarly, the number of soft-clipped bases plus mismatches may be restricted (default 3). Moreover, reads that align with a mismatch at the simulated editing site are removed. Furthermore, multi-mapped reads that align to both unedited and edited e-miRNA are considered unedited, and reads that map to several edited forms are parsimoniously attributed to the sequence(s) with the lowest number of edited sites. Each e-miRNA is then quantified based on the number of reads that align to it, with multi-mapped reads counted as *1/number of aligned e-miRNA*. Finally, all e-miRNA whose read count is below a user-defined threshold are excluded from downstream analyses (default 5). This step represents an initial identification of potentially-edited miRNA reads. Nevertheless, some reads may have originated from other transcripts, due to the alignment against e-miRBase instead of the genome or transcriptome. These will be removed in the next step.

#### Read classification

In this miRmedon step, a consensus sequence is inferred for each e-miRNA, based on its aligned clipped reads, incorporating read sequences and their Phred quality scores (**Figure 1C** and **Materials and Methods**). Consensus sequences substantially shorter or longer than the e-miRNA 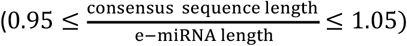 are removed from downstream analysis. Consensus sequences representing edited forms are then globally aligned to the full reference human genome with 1 mismatch using SeqMap or Bowtie, and those mapped to non-miRNA loci are discarded. Consensus sequences that do not align to the human genome are then aligned to the human transcriptome and filtered according to the same criteria. Unaligned consensus sequences and those aligned with 1 mismatch to miRNA loci or transcripts are further treated as putatively-edited miRNA sequences. Global alignment of quality-dependent consensus sequences against the genome and transcriptome allows for efficient and accurate elimination of reads that do not confidently represent edited miRNA.

#### Editing site inference and editing level quantification

In this last miRmedon step, intended to distinguish real A-to-I editing signal from sequencing errors, editing levels are modelled as a beta distribution, where the posterior editing density of each putative editing site is a beta distribution with parameters α = 1 + the number of reads with G mapped to the site, and β = 1 + the number of reads with A mapped to the site. For each putative editing site, sequencing error probabilities, computed from Phred quality scores, form an error distribution. Monte-Carlo simulation (29) is then performed to test the hypothesis that the observed substitution could be attributed to sequencing error(s). Specifically, for each putative site, random samples are repetitively drawn from the sequencing error distribution and posterior editing distribution, and are then compared. The P value, namely the probability of observing a guanosine at a primary adenosine position due to sequencing error, is calculated as the number of times in which the error rate drawn from the error distribution was greater than or equal to the editing rate drawn from the posterior editing distribution, divided by the number of resamples (**Figure 1D and Materials and Methods**). When there is no overlap between the two distributions, the P value is calculated using a binomial test, as in Alon et al. (11), with the expected probability of “success” set to the maximal value found within sequencing error distribution. Subsequently, multiple testing correction is performed using the Benjamini-Hochberg approach (22).

### miRmedon performance evaluation

To comparatively demonstrate the enhanced performance of miRmedon, we used pooled human brain small RNAseq data generated by Alon et al. for benchmarking what later became the most commonly-used miRNA editing detection tool (11). In the same pooled human brain data, miRmedon identified 32 editing sites as compared to 18 sites detected by the Alon et al. pipeline (**Supplementary Table S8**). We comprehensively annotated these sites according to previous studies. Of 129 previously reported editing sites in 98 mature miRNA, compiled by Pinto et al. (4), 75 editing sites could potentially be detected in the pooled human brain RNAseq data (based on miRNA expression in this sample). miRmedon exclusively detected five of these sites (**Figure 2A** and **Supplementary Table S8**). Moreover, we compared the detection rate of miRmedon and the Alon et al. pipeline among 58 editing sites in 55 mature miRNA that were discovered in a comprehensive survey of miRNA editing in 10,593 samples from various human tissues and conditions (4). These sites, considered high-confidence sites, were identified using the Alon et al. pipeline. Of these, in the pooled human brain data, miRmedon exclusively detected five sites (**Figure 2B** and **Supplementary Table S8**). On the other hand, three high-confidence editing sites were exclusively detected by the Alon et al. pipeline, namely hsa-miR-539-5p 10A>G, hsa-miR-376b-3p 6A>G and hsa-miR-598-3p 2 A>G. The first two were ignored by miRmedon because they were supported by reads significantly shorter than those supporting the unedited versions. The latter did not pass multiple testing correction. The editing levels of these sites are not significantly lower than other sites, ruling out the possibility that detection threshold differences could have accounted for the increased sensitivity demonstrated by miRmedon (**Figure 2C**). Moreover, several editing sites exclusively predicted by Alon et al. were found to overlap with gnomAD variants (**Supplementary Table S3** and **Supplementary Table S4**), which, by definition, are not predicted by miRmedon, thereby demonstrating its improved specificity.

**Figure 2.**
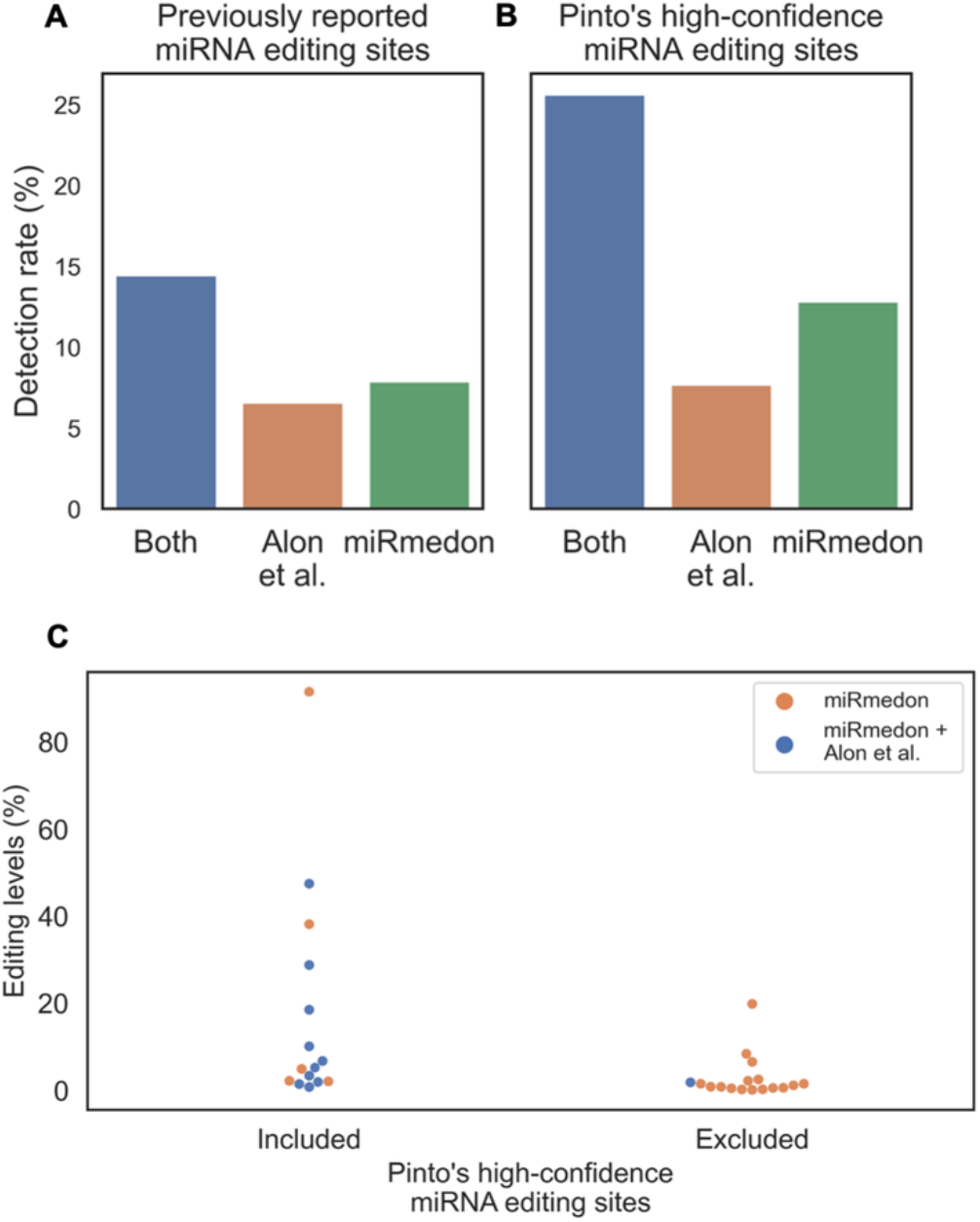
Improved sensitivity of miRmedon as compared to the most commonly used miRNA editing detection tool developed by Alon et al. (11). Head to head comparison of miRmedon and the Alon et al. pipeline among (**A**) all previously reported editing sites compiled by Pinto et al., and (**B**) the subset of editing sites considered to be of high-confidence by Pinto et al. (4). The detection rate was calculated as the number of editing sites reported by Alon et al. and miRmedon in small RNAseq from pooled human brain tissue, ignoring putative editing sites that overlap gnomAD SNPs (Supplementary Table S3) or those within 5bp of gnomAD indels (Supplementary Table S4), and considering miRNA expression in the analyzed sample; (**C**) Among Pinto’s high-confidence sites, five sites were exclusively detected by miRmedon, independent of their editing levels (Supplementary Table S8).

We next estimated miRmedon’s signal-to-noise ratio for A-to-I miRNA editing detection, by quantifying the proportion of A>G substitutions relative to all other possible substitutions. Toward that end, we generated e-miRBase indices for all 12 possible nucleotide substitutions and processed the pooled human brain small RNAseq reads according to the miRmedon pipeline for each substitution. A>G substitutions represented 97.8% of all miRmedon-detected substitutions, while every other substitution accounted for <1% (**Figure 3**).

**Figure 3.**
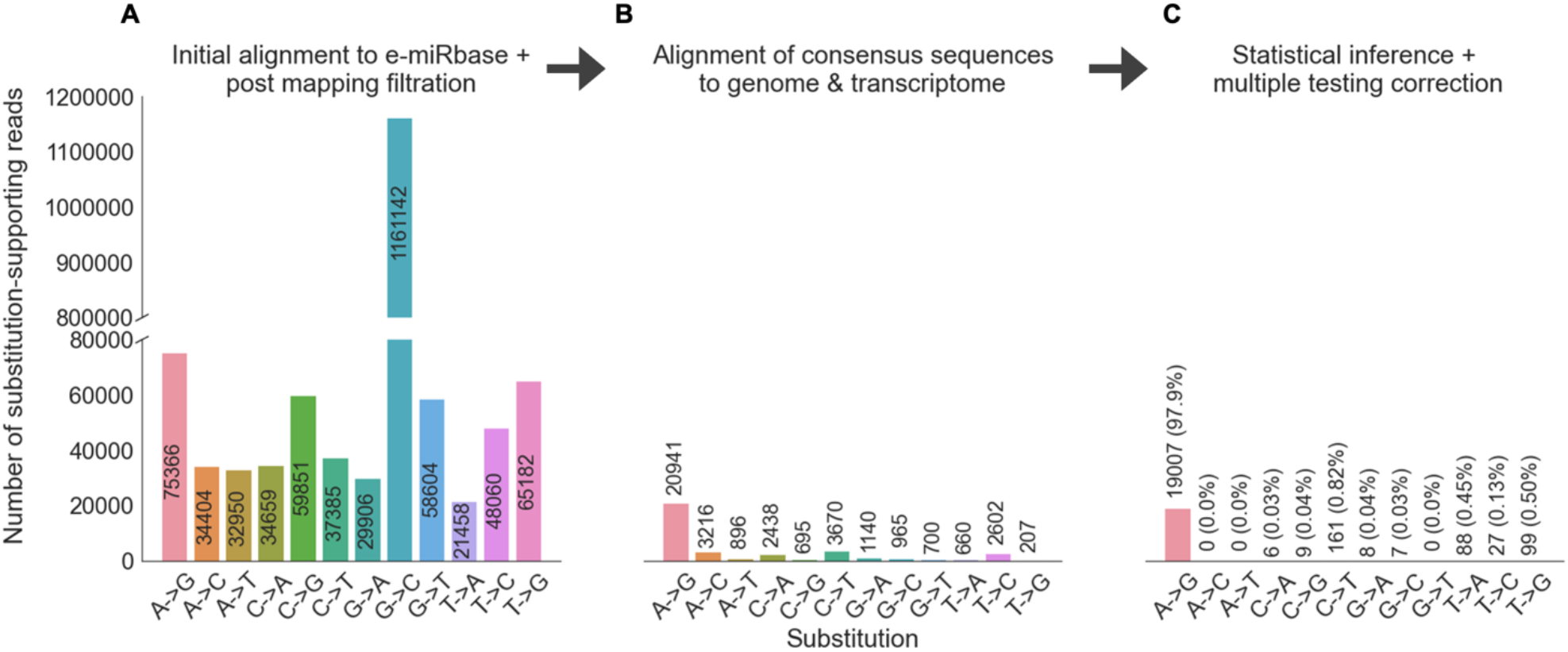
Serial increase in specificity offered by each miRmedon step. Shown is the total number of substitution-supporting reads for each of the 12 possible substitutions in pooled human brain small RNAseq data, after each main miRmedon step. While the total number of substitution-supporting reads decreases with each step, a strong signal of A>G substitutions remains, thereby demonstrating miRmedon’s high specificity for A-to-I miRNA editing sites. (**A**) Number or reads supporting each substitution following local alignment to e-miRBase and post mapping filtration; (**B**) Number of reads supporting each substitution following global alignment of consensus sequences to the human genome and transcriptome; (**C**) Number of reads supporting each substitution after statistical inference and multiple testing correction.

The miRmedon pipeline includes two major filtration steps intended to address two major sources of possible false positives, namely (1) misclassifying other small RNAs or RNA fragments as edited miRNA, and (2) misidentifying editing due to sequencing errors. We therefore compared the number of reads supporting each substitution with respect to these steps. An inflation of all possible 12 substitutions was observed following initial alignment of pooled human brain small RNAseq reads against the 12 e-miRBases and post alignment filtration steps, which include removal of reverse complement alignments, removal of reads with more than two soft-clipped bases, and removal of reads that align with a mismatch at the simulated editing site (**Figure 3A**). Of note, potential false positives arising from (1) the misidentification of editing due to a genomic variant at the site or its vicinity, and (2) the highly modified 3’ ends of miRNA, are handled during the e-miRBase generation phase of miRmedon. Therefore, the counts depicted in Figure 3A are independent of these sources of noise. It should also be noted that the number of reads supporting G>C substitutions after initial alignment to e-miRBase, and following post-alignment filters, was 15.4-fold higher than that of the A>G ones (**Figure 3B**). miRmedon’s approach to tackle this issue by integrating all e-miRNA supporting reads into quality-dependent consensus sequences and aligning these to the genome and transcriptome shows prominent efficacy in reducing this type of false positive results, seeing that following this step the only substantial signal was that of A>G substitutions (**Figure 3C**). Specifically, 965 substitutions were retained in this step, supported by 38,130 reads, of which A>G ones represented 54.9%. Then, in addressing the misidentification of sequencing errors as editing sites, miRmedon’s original statistical inference approach substantially improved the signal of A>G substitutions relative to all other types of mismatches (**Figure 3C**). Specifically, following multiple testing correction, a clear signal of A>G substitutions emerged, supported by 18,853 reads and representing 97.8% of all possible substitutions. See **Supplementary Table S9** for their distribution.

That 97.8% of mismatch-supporting reads support A>G mismatches out of all possible mismatches is consistent with ADAR enzymes being the dominant editing enzymes in the human brain. Yet, ADARs are not the only active RNA editors in the human brain. Activation induced cytidine deaminases/apolipoprotein B mRNA editing enzyme cytidine deaminases (AID/APOBECs) are lowly expressed in the human brain (30), and C-to-U editing of miRNA has been reproducibly reported in both molecular and computational screens (11,31-36). Moreover, recent applications of the Alon et al. pipeline revealed recurrent non-canonical miRNA editing in human brain tissue (32,33). At the site level, miRmedon identified 32 A>G sites, 16 lowly edited C>T sites, and 10 non-canonical mismatches in the pooled human brain sample (**Supplementary Table S6**). In other words, its specificity for canonical editing is 82.759%.

We next turned to assess miRmedon’s false discovery rate (FDR). With a goal of estimating miRmedon’s FDR using mismatches detected in the pooled human brain RNA sample, we asked whether we could identify miRmedon-detected sites that most closely resemble high-confidence A-to-I miRNA editing sites compiled by Pinto et al. (4), and in doing so identify sites that are significantly different from such high-confidence sites – these could be considered false positives. We accordingly trained a regularized logistic regression model to distinguish between high-confidence miRNA editing sites and all other sites (**Supplementary Table S5**). The model, which is based on miRmedon-reported metrics as well as the sequence context of each site and whether it is located in the miRNA seed region, performs with an area under the receiver operating characteristics (ROC) curve (AUC) of 0.84 (**Figure 4A**). Thus, despite the small sample size this model is quite accurate. When applied to all miRmedon-detected sites, using a prediction probability cutoff of 0.3, the model classified 31 of 32 A>G sites as similar to high-confidence *bona fide* A-to-I miRNA editing sites (**Figure 4B** and **Supplementary Table S7**). That is, the model estimates miRmedon’s FDR to be 1/32 = 3.125%. Specifically, the model flags the A>G mismatch in position 8 of hsa-miR-376c-3p as a false positive. This site had the largest adjusted P value (0.032), which is essentially correlated to the FDR expectation. A conservative estimation of miRmedon’s FDR, considering the model’s results and specificity for canonical editing, would be between 3.125% and 17.241%.

**Figure 4.**
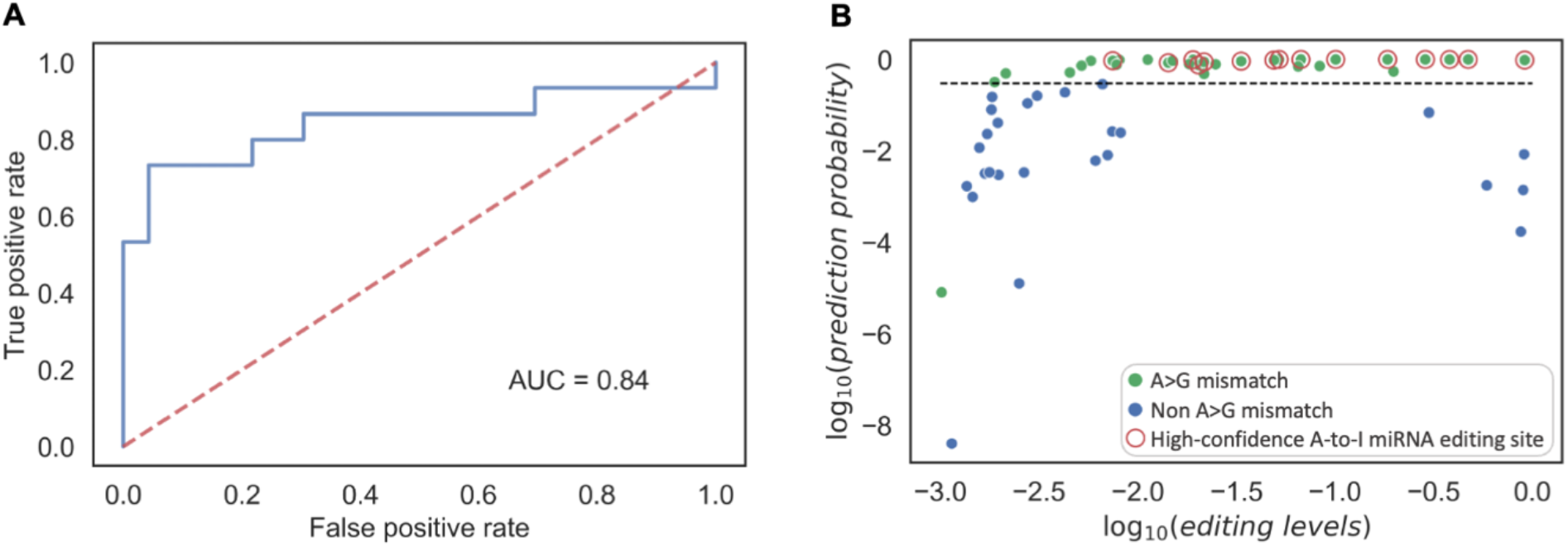
FDR estimation according to similarity to high-confidence A-to-I miRNA editing sites. (**A**) ROC curve of a multivariate model trained to distinguish high-confidence A-to-I miRNA editing sites from non A>G substitutions. See Supplementary Table S5 for model features and their importance. ROC computed using leave one out cross validation; (**B**) Application of the model to all 58 mismatches detected in pooled human brain RNA. Shown is a log-log plot of the model’s prediction probability as a function of the editing level. A>G sites are marked in green, non A>G sites are in blue, and Pinto’s high-confidence A-to-I miRNA editing sites are circled in red. A threshold of -0.523, which corresponds to a probability of 0.3 is shown by a dashed black line. A clear separation can be seen between most A>G sites and non A>G sites. According to the model, 31 of 32 A>G substitutions closely resemble high-confidence A-to-I miRNA editing sites (at a prediction probability cutoff of 0.3). Thus the model estimates miRmedon’s FDR to be 1/32 (3.125%).

### Validation

We used two independent validation approaches. The first followed the Alon et al. strategy for editing site validation using ADAR or ADARB1 overexpression in glioblastoma cell lines. Specifically, for each site originally detected in the pooled human brain small RNAseq data, overediting in the cell lines overexpressing ADAR or ADARB1 as compared to their wildtype counterparts was considered as successful validation. Following this strategy, eight editing sites detected by miRmedon were validated in U118 cells stably overexpressing ADARB1, as compared to control U118 cells (**Figure 5**). Of these, only two sites (hsa-miR-455 17A>G and hsa-miR-421 14A>G) were also reported (and validated) by Alon et al. (2012). In addition, two validated A-to-I editing sites (hsa-miR-365a\b-3p 11 A>G and hsa-miR-140-5p 16A>G) are first reported in this work. No miRmedon-detected sites were validated in ADAR-overexpressing cells. While Alon et al. validated ADAR-mediated editing of the fourth position of hsa-miR-381-3p, this miRNA was excluded from miRmedon analyses due to low coverage (four reads, while the threshold was set to five reads).

**Figure 5.**
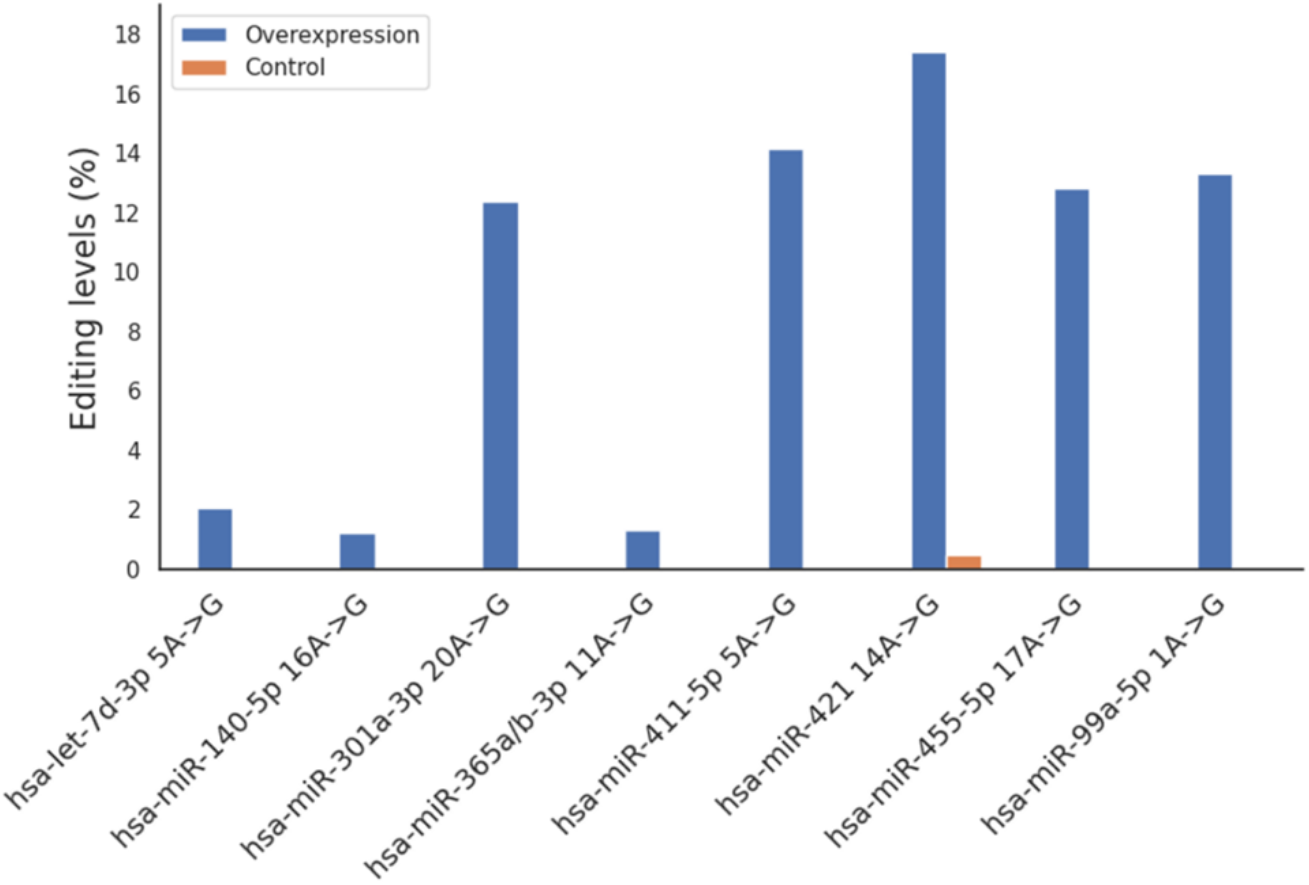
Eight editing sites validated by stable overexpression of ADARB1 in U118 glioblastoma cell line. All eight sites, originally detected in a pooled human brain sample, were also detected in U118 overexpressing ADARB1 but not in U118 control cells. Two of these, hsa-miR-365a/b 11A>G and hsa-miR-140-5p 16A>G, have not been previously reported to be A-to-I miRNA editing sites.

As a second validation, we used 64 small RNAseq samples of postmortem prefrontal cortex from individuals with Huntington’s Disease and neurotypical individuals (GSE64977) (37). For each site originally detected in the pooled human brain RNA, editing detected in three or more postmortem cortical samples was considered successful validation. Of 32 sites detected by miRmedon in the pooled human brain RNA, 26 were validated in this independent multi-sample dataset, one was not validated, and the remaining five sites could not be validated because their host miRNA was expressed below detection limits (**Supplementary Table S8**). Thus, based on this independent validation, miRmedon’s FDR could be estimated to be 1/27 = 3.70%. This estimate is within the FDR range described above ([3.125%, 17.241%]).

### Characteristics of newly detected sites

We compared the characteristics of 15 sites first reported in this work to 17 previously reported sites detected in the same small RNAseq data (**Supplementary Table S8**). By and large, currently available approaches, including that of Alon et al., cannot detect editing sites in multi-mapped miRNAs and sites observed only in the vicinity of another site. miRmedon exclusively detected seven A-to-I editing sites within multi-mapped miRNAs (hsa-miR-365a/b-3p and hsa-miR-376a-3p), and six sites found only as part of 2-site haplotypes (five hsa-miR-376a-3p haplotypes and one hsa-miR-376c-3p haplotype). Moreover, of the 15 sites that were exclusively detected by miRmedon, six reside within the seed region, with the potential of altering miRNA targeting. Additionally, owing to its novel statistical inference approach, miRmedon is capable of confidently detecting lowly edited sites, and seven such sites were detected here for the first time. Finally, 10 miRmedon-specific sites were detected in three or more postmortem prefrontal cortex samples of GSE64977, and the remaining five miRmedon-specific sites were in miRNA whose expression levels were below detection limit in these cortical samples (**Supplementary Table S8**).

### Detection of co-edited mature miRNA

One unique feature of miRmedon, as compared to all other existing tools for miRNA editing detection, is its ability to detect haplotypes of co-editing at an unlimited number of sites. Six 2-site haplotypes were identified by miRmedon in pooled human brain data: three haplotypes of hsa-miR-376a-3p, two of hsa-miR-376c-3p and one of hsa-miR-381-3p (**Supplementary Table S9**). Of these, editing at the fourth and seventh positions of hsa-miR-381-3p, each deemed by Pinto et al. to be a high-confidence editing site, were co-detected for the first time, in 32 reads. The predicted secondary structure of hsa-miR-381 remained stable in hairpin form even with inosine at both editing sites (**Figure 6**). Moreover, haplotypes of the hsa-miR-376 family were found to contain a dominant editing site at the sixth position, deemed by Pinto et al. to be a high-confidence site, co-occurring with editing at the 8^th^, 12^th^, 13^th^, 14^th^, or 18^th^ position. The present study is the first to report these haplotypes.

**Figure 6.**
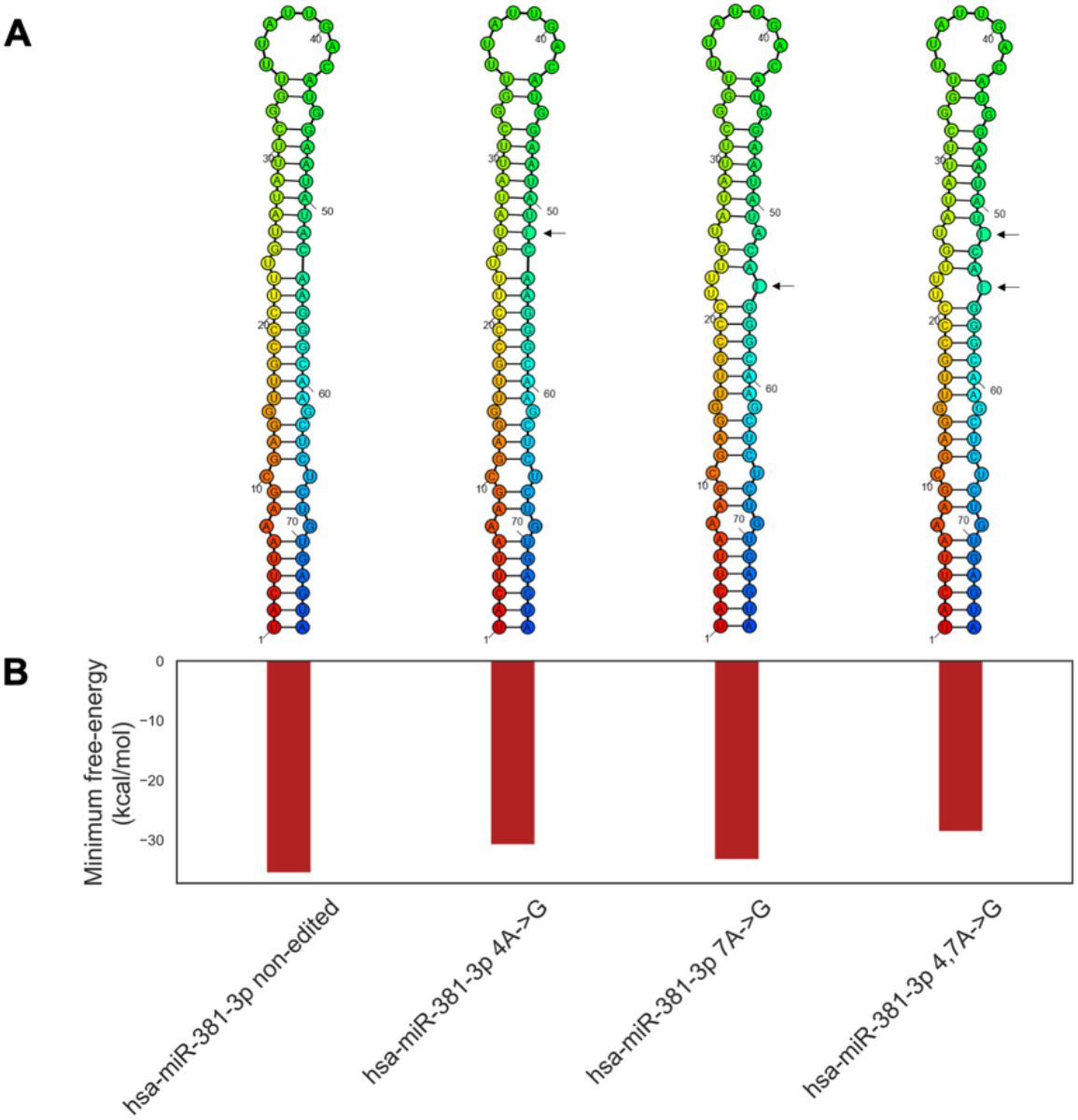
Multiple editing sites detected by miRmedon in hsa-miR-381-3p (**Supplementary Table S9** and **S12**). Two editing sites at the fourth and seventh positions (indicated by arrows) were deemed to be high-confidence miRNA editing sites by Pinto et al. (4), although Alon et al. (11) reported only on the first. miRmedon identified both sites as significant A-to-I miRNA editing sites, with 9,988 reads supporting the co-occurrence of ‘G’ at both positions (i.e. in the same read) in 54 samples. (**A**) The predicted secondary structure of hsa-miR-381 pre-miRNA remained stable with the presence of inosine at either or both editing sites; (**B**) the estimated minimum free-energy of the pre-miRNA hairpin remained around -30kcal/mol.

Finally, we used miRmedon to examine the scope and characteristics of mature miRNA co-editing in human prefrontal cortex, using small RNAseq data from 64 postmortem tissue samples of Broadman Area 9, obtained from individuals with Huntington’s disease and neurotypical controls (GSE64977, Ref. 37). miRmedon detected nine doubly edited mature miRNA forms (**Figure 7**), each found in at least three samples, and supported by 99.22 reads, on average (**Supplementary Tables S10-12**). Eight of the nine doubly edited mature miRNA detected in human prefrontal cortex included an edited site within the miRNA seed region, potentially also affecting targeting. The most common editing haplotype was that of positions 20 and 21 in hsa-miR-301b-3p, co-edited in all cortical samples. This doubly edited hsa-miR-301b-3p form was more common than hsa-miR-301b-3p edited in position 21 alone. Furthermore, two doubly edited forms of hsa-miR-381-3p included two miRNA seed sites, with the potential of effectively shifting this miRNA’s targeting: Co-editing of hsa-miR-381a-3p positions 4 and 7 was found in 83% of samples, with a subset of these samples also expressing a doubly edited hsa-miR-381a-3p at positions 6 and 7. Both of these co-edited forms of hsa-miR-381-3p were also detected by miRmedon in the Alon et al. pooled human brain data.

**Figure 7.**
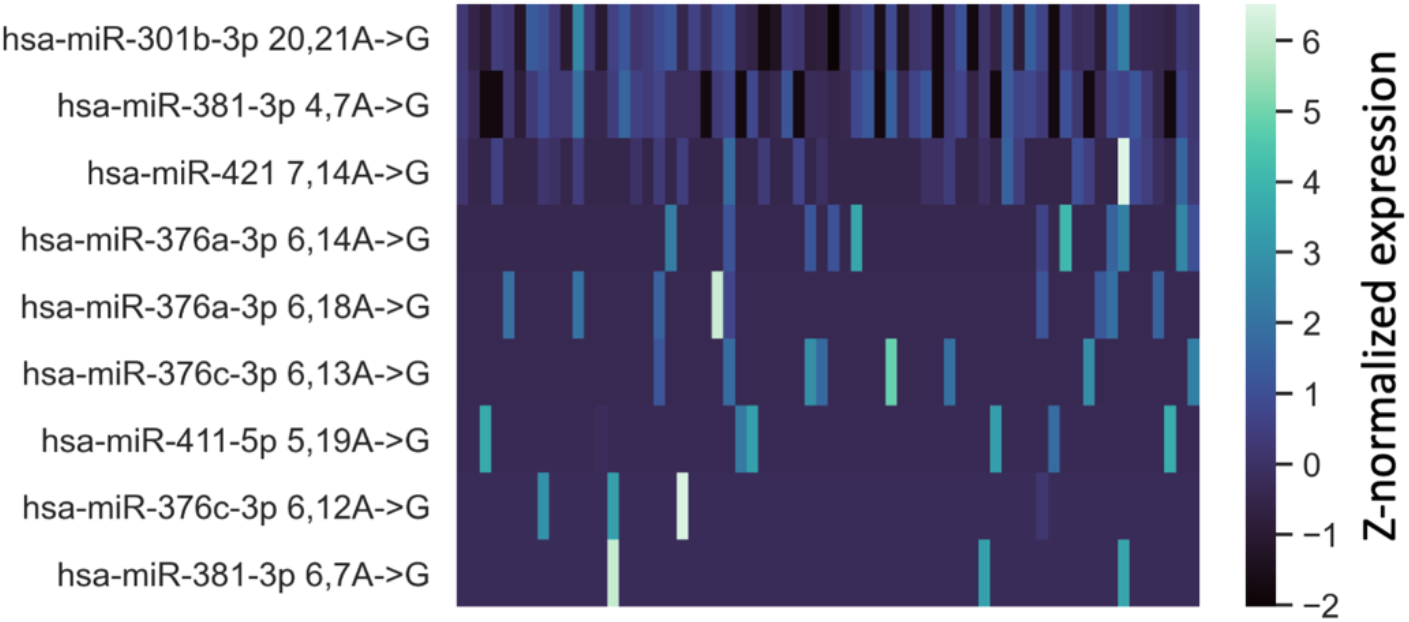
Doubly edited mature miRNA detected by miRmedon in 64 human prefrontal cortex samples. Shown are the normalized CPM value of nine doubly edited miRNA haplotypes detected in at least three samples, and supported by at least five reads in each sample. Three 2-site editing haplotypes, namely hsa-miR-381-3p 4,7A->G, hsa-miR-376a-3p 6,14A->G, and hsa-miR-376a-3p 6,18A->G, were also detected in an independent pooled human brain sample.

Moreover, hsa-miR-376a-3p was edited at both positions 6 and 14, or positions 6 and 18, but not in any one site alone. Editing of its close relative hsa-miR-376c-3p was detected in position 6 in all samples, and a small fraction of these expressed doubly edited forms of hsa-miR-376c-3p at positions 6 and 12, and/or positions 6 and 13. Similarly, hsa-miR-411-5p was edited in position 5 in all samples, with 11% also expressing a doubly-edited form, with “G” at positions 5 and 19. Finally, hsa-miR-421 was edited in position 7 and position 14, with co-editing found only in those samples expressing both forms of single-site (position 7 or position 14) hsa-miR-421 editing.

## DISCUSSION

### Advantages over existing methods

#### Improved sensitivity

By considering all possible combinations of ‘A’ to ‘G’ substitutions at non-variance-prone adenosine sites, miRmedon enables the discovery of multiple editing events per read. As a result, miRmedon facilitates the investigation of miRNA editing linkage, for the first time. Further improvement in sensitivity over existing methods arises from miRmedon’s unconventional approach that goes beyond focusing only on mismatches to identify overrepresented ones, thereby enabling the identification of edited miRNA harboring a SNP in cis to the editing site.

Another key improvement in sensitivity offered by miRmedon addresses the problem of overrepresented RNA modifications at the 3’ end of mature miRNA (38), which dampens the sensitivity of global aligners, such as Bowtie, that have been utilized in a number of existing methods (**Supplementary Table S1**). In addition, widely used Illumina sequencing often demonstrates low quality calls at the read ends, further contributing to this predicament. To overcome these challenges, miRmedon uses the local alignment mode of STAR, which allows for soft clipping, to align small RNAseq reads to e-miRBase. This functionality spares the suggested transversal trimming of bases at the 3’ end (11), thereby improving alignment specificity.

Furthermore, the statistical inference employed by miRmedon for distinguishing an A-to-I editing signal from sequencing error enables the confident detection of lowly edited sites. Finally, by considering all human miRNA irrespective of the number of their genomic loci, miRmedon is the only currently available tool capable of detecting editing of the 1% of human miRNA found in > 100 loci in the human genome (**Supplementary Table S1**).

#### Improved specificity

miRmedon leverages large scale population variation data not only for improved sensitivity (detailed above), but also for ensuring that the observed A/G mixture is not a result of a rare SNP or misalignment due to a nearby indel. **Supplementary Tables S3** and **S4** detail previously reported miRNA editing sites that overlap with genomic SNPs or indels, respectively, as reported by gnomAD.

Moreover, the cross-mapping challenge (6), which is aggravated by considering all possible combinations of putative editing sites, is addressed by globally aligning a data-driven consensus sequence for each e-miRNA to the genome using aligners capable of finding all possible loci from which a read could have originated (e.g. SeqMap or Bowtie). Moreover, the use of the full human reference genome (including scaffolds, assembly patches, and haplotypes) further contributes to enhanced specificity. Finally, additional specificity is gained by aligning consensus sequences to the transcriptome so as to exclude the possibility of a spliced RNA fragment being considered as edited miRNA.

The miRmedon output could be filtered depending on the desired sensitivity-specificity trade-off, for example by considering editing supported by a number of reads above a certain threshold, sites detected in multiple samples, or any combination of filters suggested by Pinto and Levanon (39).

### Limitations

Like other existing methods, miRmedon cannot confidently detect editing at the 3’ ends of miRNA. Furthermore, it does not detect editing in SNPs, in close proximity to indels, or when edited miRNA perfectly match another genomic location. Finally, its accuracy is relatively low for editing sites supported by a few reads.

### Co-editing of mature miRNA in the adult human brain

Although miRmedon can confidently detect co-editing of up to 12 sites in mature miRNA of miRBase 21, only doubly edited mature miRNA were detected. This finding suggests that hyper-editing of miRNA is not common in the adult human prefrontal cortex.

## DATA AVAILABILITY

miRmedon is available as an open source tool at https://github.com/Amitai88/miRmedon.

All sequencing data processed as part of this study is available through the to the NCBI Gene Expression Omnibus (https://www.ncbi.nlm.nih.gov/geo/), via accession numbers detailed in Supplementary Table S2.

## SUPPLEMENTARY DATA

Supplementary Data are available at NAR online.

## ACKNOWLEDGEMENTS

We thank Jerry Eichler and members of the Eran lab for their great help.

## FUNDING

This research was supported by the Israel Science Foundation [2755/20 to A.E.]. A.M. was supported by the Israeli Scholarship Education Foundation (ISEF).

## CONFLICT OF INTEREST

The authors declare no conflict of interest.

## Notes

### Competing Interest Statement

The authors have declared no competing interest.

## REFERENCES

1. Tassinari, V., Cesarini, V., Silvestris, D.A. and Gallo, A. (2019) The adaptive potential of RNA editing-mediated miRNA-retargeting in cancer. Biochim Biophys Acta Gene Regul Mech, 1862, 291–300.

2. Behm, M. and Ohman, M. (2016) RNA Editing: A Contributor to Neuronal Dynamics in the Mammalian Brain. Trends Genet, 32, 165–175.

3. Wang, Y. and Liang, H. (2018) When MicroRNAs Meet RNA Editing in Cancer: A Nucleotide Change Can Make a Difference. Bioessays, 40.

4. Pinto, Y., Buchumenski, I., Levanon, E.Y. and Eisenberg, E. (2018) Human cancer tissues exhibit reduced A-to-I editing of miRNAs coupled with elevated editing of their targets. Nucleic Acids Res, 46, 71–82.

5. Landgraf, P., Rusu, M., Sheridan, R., Sewer, A., Iovino, N., Aravin, A., Pfeffer, S., Rice, A., Kamphorst, A.O., Landthaler, M. et al. (2007) A mammalian microRNA expression atlas based on small RNA library sequencing. Cell, 129, 1401–1414.

6. de Hoon, M.J., Taft, R.J., Hashimoto, T., Kanamori-Katayama, M., Kawaji, H., Kawano, M., Kishima, M., Lassmann, T., Faulkner, G.J., Mattick, J.S. et al. (2010) Cross-mapping and the identification of editing sites in mature microRNAs in high-throughput sequencing libraries. Genome Res, 20, 257–264.

7. Pantano, L., Estivill, X. and Marti, E. (2010) SeqBuster, a bioinformatic tool for the processing and analysis of small RNAs datasets, reveals ubiquitous miRNA modifications in human embryonic cells. Nucleic Acids Res, 38, e34.

8. Vesely, C., Tauber, S., Sedlazeck, F.J., von Haeseler, A. and Jantsch, M.F. (2012) Adenosine deaminases that act on RNA induce reproducible changes in abundance and sequence of embryonic miRNAs. Genome Res, 22, 1468–1476.

9. Ekdahl, Y., Farahani, H.S., Behm, M., Lagergren, J. and Ohman, M. (2012) A-to-I editing of microRNAs in the mammalian brain increases during development. Genome Res, 22, 1477–1487.

10. Zheng, Y., Ji, B., Song, R., Wang, S., Li, T., Zhang, X., Chen, K., Li, T. and Li, J. (2016) Accurate detection for a wide range of mutation and editing sites of microRNAs from small RNA high-throughput sequencing profiles. Nucleic Acids Res, 44, e123.

11. Alon, S., Mor, E., Vigneault, F., Church, G.M., Locatelli, F., Galeano, F., Gallo, A., Shomron, N. and Eisenberg, E. (2012) Systematic identification of edited microRNAs in the human brain. Genome Res, 22, 1533–1540.

12. Griffiths-Jones, S., Grocock, R.J., van Dongen, S., Bateman, A. and Enright, A.J. (2006) miRBase: microRNA sequences, targets and gene nomenclature. Nucleic Acids Res, 34, D140–144.

13. Musumeci, L., Arthur, J.W., Cheung, F.S., Hoque, A., Lippman, S. and Reichardt, J.K. (2010) Single nucleotide differences (SNDs) in the dbSNP database may lead to errors in genotyping and haplotyping studies. Hum Mutat, 31, 67–73.

14. Picardi, E. and Pesole, G. (2013) REDItools: high-throughput RNA editing detection made easy. Bioinformatics, 29, 1813–1814.

15. John, D., Weirick, T., Dimmeler, S. and Uchida, S. (2017) RNAEditor: easy detection of RNA editing events and the introduction of editing islands. Brief Bioinform, 18, 993–1001.

16. Langmead, B. (2010) Aligning short sequencing reads with Bowtie. Curr Protoc Bioinformatics, Chapter 11, Unit 11 17.

17. Jiang, H. and Wong, W.H. (2008) SeqMap: mapping massive amount of oligonucleotides to the genome. Bioinformatics, 24, 2395–2396.

18. Dobin, A., Davis, C.A., Schlesinger, F., Drenkow, J., Zaleski, C., Jha, S., Batut, P., Chaisson, M. and Gingeras, T.R. (2013) STAR: ultrafast universal RNA-seq aligner. Bioinformatics, 29, 15–21.

19. Ziemann, M., Kaspi, A. and El-Osta, A. (2016) Evaluation of microRNA alignment techniques. RNA, 22, 1120–1138.

20. Katoh, K., Misawa, K., Kuma, K. and Miyata, T. (2002) MAFFT: a novel method for rapid multiple sequence alignment based on fast Fourier transform. Nucleic Acids Res, 30, 3059–3066.

21. Katoh, K. and Standley, D.M. (2013) MAFFT multiple sequence alignment software version 7: improvements in performance and usability. Mol Biol Evol, 30, 772–780.

22. Benjamini, Y. and Hochberg, Y. (1995) Controlling the False Discovery Rate: A Practical and Powerful Approach to Multiple Testing. Journal of the Royal Statistical Society. Series B (Methodological), 57, 289–300.

23. Andrews, S.,. Available online at: http://www.bioinformatics.babraham.ac.uk/projects/fastqc. (2010).

24. Martin, M. (2011) Cutadapt removes adapter sequences from high-throughput sequencing reads. 2011, 17, 3.

25. Bolger, A.M., Lohse, M. and Usadel, B. (2014) Trimmomatic: a flexible trimmer for Illumina sequence data. Bioinformatics, 30, 2114–2120.

26. Karczewski, K.J., Francioli, L.C., Tiao, G., Cummings, B.B., Alföldi, J., Wang, Q., Collins, R.L., Laricchia, K.M., Ganna, A., Birnbaum, D.P. et al. (2020) The mutational constraint spectrum quantified from variation in 141,456 humans. Nature, 581, 434–443.

27. Lorenz, R., Bernhart, S.H., Honer Zu Siederdissen, C., Tafer, H., Flamm, C., Stadler, P.F. and Hofacker, I.L. (2011) ViennaRNA Package 2.0. Algorithms Mol Biol, 6, 26.

28. Lu, J.S., Bindewald, E., Kasprzak, W.K. and Shapiro, B.A. (2018) RiboSketch: versatile visualization of multi-stranded RNA and DNA secondary structure. Bioinformatics, 34, 4297–4299.

29. North, B.V., Curtis, D. and Sham, P.C. (2002) A note on the calculation of empirical P values from Monte Carlo procedures. Am J Hum Genet, 71, 439–441.

30. Correia de Sousa, M., Gjorgjieva, M., Dolicka, D., Sobolewski, C. and Foti, M. (2019) Deciphering miRNAs’ Action through miRNA Editing. Int J Mol Sci, 20.

31. Negi, V., Paul, D., Das, S., Bajpai, P., Singh, S., Mukhopadhyay, A., Agrawal, A. and Ghosh, B. (2015) Altered expression and editing of miRNA-100 regulates iTreg differentiation. Nucleic Acids Res, 43, 8057–8065.

32. Paul, D., Ansari, A.H., Lal, M. and Mukhopadhyay, A. (2020) Human Brain Shows Recurrent Non-Canonical MicroRNA Editing Events Enriched for Seed Sequence with Possible Functional Consequence. Noncoding RNA, 6.

33. Paul, D., Sinha, A.N., Ray, A., Lal, M., Nayak, S., Sharma, A., Mehani, B., Mukherjee, D., Laddha, S.V., Suri, A. et al. (2017) A-to-I editing in human miRNAs is enriched in seed sequence, influenced by sequence contexts and significantly hypoedited in glioblastoma multiforme. Sci Rep, 7, 2466.

34. Wang, Q., Zhao, Z., Zhang, X., Lu, C., Ren, S., Li, S., Guo, J., Liao, P., Jiang, B. and Zheng, Y. (2019) Identifying microRNAs and Their Editing Sites in Macaca mulatta. Cells, 8.

35. Gong, J., Wu, Y., Zhang, X., Liao, Y., Sibanda, V.L., Liu, W. and Guo, A.Y. (2014) Comprehensive analysis of human small RNA sequencing data provides insights into expression profiles and miRNA editing. RNA Biol, 11, 1375–1385.

36. Guo, S., Yang, J., Jiang, B., Zhou, N., Ding, H., Zhou, G., Wu, S., Suo, A., Wu, X., Xie, W. et al. (2022) MicroRNA editing patterns in Huntington’s disease. Sci Rep, 12, 3173.

37. Hoss, A.G., Labadorf, A., Latourelle, J.C., Kartha, V.K., Hadzi, T.C., Gusella, J.F., MacDonald, M.E., Chen, J.F., Akbarian, S., Weng, Z. et al. (2015) miR-10b-5p expression in Huntington’s disease brain relates to age of onset and the extent of striatal involvement. BMC Med Genomics, 8, 10.

38. Burroughs, A.M., Ando, Y., de Hoon, M.J., Tomaru, Y., Nishibu, T., Ukekawa, R., Funakoshi, T., Kurokawa, T., Suzuki, H., Hayashizaki, Y. et al. (2010) A comprehensive survey of 3’ animal miRNA modification events and a possible role for 3’ adenylation in modulating miRNA targeting effectiveness. Genome Res, 20, 1398–1410.

39. Pinto, Y. and Levanon, E.Y. (2019) Computational approaches for detection and quantification of A-to-I RNA-editing. Methods, 156, 25–31.

